# Ozone Treatment for Elimination of Bacteria and SARS-CoV-2 for Medical Environments

**DOI:** 10.1101/420737

**Authors:** Craig Westover, Savlatjon Rahmatulloev, David Danko, Ebrahim Afshinnekoo, Niamh B. O’Hara, Rachid Ounit, Daniela Bezdan, Christopher E. Mason

**Affiliations:** Department of Physiology and Biophysics, Weill Cornell Medical College, New York, NY, USA; The HRH Prince Alwaleed Bin Talal Bin Abdulaziz Alsaud Institute for Computational Biomedicine, Weill Cornell Medical College, New York, NY, USA; The WorldQuant Initiative for Quantitative Prediction, Weill Cornell Medicine, New York, NY, USA; Jacobs Technion-Cornell Institute, Cornell Tech, New York, NY, USA; The Feil Family Brain and Mind Research Institute, New York, NY, USA

**Keywords:** Microbiome, SARS-CoV-2, HAI, CFU, ozone, decontamination, log-kill, built-environment, RNA degradation

## Abstract

Pathogenic bacteria and viruses in medical environments can lead to treatment complications and hospital-acquired infections (HAIs), and current cleaning protocols do not address hard-to-access areas or that may be beyond line-of-sight treatment such as with ultraviolet radiation. At the time of writing, the ongoing pandemic of the novel coronavirus known as novel coronavirus (2019-nCoV) has claimed over 4 million cases worldwide and is expected to have multiple peaks, with possible resurgences throughout 2020. It is therefore imperative that disinfection methods in the meantime be employed to keep up with the supply of personal protective equipment (PPE) and sterilize a wide array of surfaces as quarantine lockdowns begin to be lifted.

Here, we tested the efficacy of Sani Sport ozone devices as a means to treat hospital equipment and surfaces for killing bacteria, degrading synthetic severe acute respiratory syndrome coronavirus 2 (SARS-CoV-2) RNA, and RNA from non-replicative capsid enclosed SARS-CoV-2. We observed a rapid killing of medically-relevant and environmental bacteria (*Escherichia coli*, *Enterococcus faecalis*, *Bacillus subtlis*, and *Deinococcus radiodurans*) across four surfaces (blankets, catheter, remotes, and syringes) within 30 minutes, and up to a 99% reduction in viable bacteria at the end of 2-hour treatment cycles. Significant RNA degradation of synthetic SARS-CoV-2 RNA was seen an hour into the ozone treatment as compared to non-treated controls and a non-replicative form of the virus was shown to have significant RNA degradation at 30 minutes compared to a no treatment control and RNA degradation could be reliably detected at 10,000 and 1,000 copies of virus per sample. These results show the strong promise of ozone treatment for reducing risk of infection and HAIs.

## Introduction

The World Health Organization (WHO) has recently stated that we are entering a “postantibiotic era” due to decades of overuse of antibiotics for therapeutic and agricultural reasons^1^. Antimicrobial resistance (AMR) has been increasing at an alarming rate, whereby prevalent nosocomial infections such as pneumonia, tuberculosis, methicillin-resistant *Staphylococcus aureus* (MRSA), and *Clostridium difficile* infection are becoming difficult to treat with traditional methods, due to multidrug resistance (MDR) and AMR. In the United States alone, hospital acquired infections (HAIs) kill an average of 63,000 patients yearly, with many showing AMR. In order to combat this crisis, resources have been allocated to developing modified versions of existing antibiotics or discovery of new ones. However, resistance to new and existing drugs continues to persist due to the strong selective pressure of antibiotics, with successful bacteria acquiring antimicrobial resistance genes in a continual ‘arms race’ between antibiotic development and antibiotic resistance^,3^.

As such, there is a need of novel approaches for effective disinfection tools that control drug resistant pathogens and reduce antibiotic utilization and consumption. In the past, ozone has been utilized to safely sanitize a number of products in various industries, such as sewage treatment to kill harmful bacteria^4^. The strong electronegative properties of ozone encourages disruption of proteins, peptidoglycans, and lipids in the cell wall and cell membrane, and interferes with the activity of enzymes and nucleic acids. It has been demonstrated that ozone is up to 3,000 times faster acting and 150 times stronger than chlorine for killing bacteria, fungi, and other pathogens under some conditions^5^. Also, ozone has been used at highly toxic concentrations to sanitize hospital rooms prior to patient occupation^6,7^. However, ozone is inherently unstable as it degrades to diatomic oxygen if its source of production is exhausted^5^. Thus, a method for continually-generating ozone may be an ideal tool for sanitizing various hospital surfaces or equipment, as an alternative to the autoclave or ultraviolet light, but such a device (to our knowledge) has not been tested on medical equipment. Here, we report the impact of an ozone-generating machine on medically-relevant surfaces and materials from a hospital, to gauge the impact on bacterial growth (colony forming units, or CFUs) and response (percent of bacteria killed).

Ozone disinfection of surfaces and hospital personal protective equipment (PPE) has been suggested to be a more eco-friendly disinfectant with a short half-life, leaving no chemical byproducts behind. It has been demonstrated that standard N95 mask filters can withstand many cycles of ozone gas treatment with no degradation of material or filtration efficiency even up to 200 part per million (ppm) for 90 minutes or 20 ppm for up to 36 hours, both dosages well beyond the proposed viricidal dose of 10 ppm for 30 minutes necessary for 99% viral inactivation^8,9^. In addition, gaseous ozone has the benefit of being able to spread throughout a room and was shown to penetrate multiple crevices and surfaces compared to liquid sprays or aerosols in order to kill different types of viruses including aerosol-borne viruses^10^.

2019-nCoV is an enveloped RNA virus which makes it more susceptible to ozone degradation than naked virions as ozone is known to interact with envelope and capsid proteins via formation of protein hydroxides and protein hydroperoxides^11^. Unsaturated lipids, carbohydrates, and nucleic acids are also damaged by ozone with damage to nucleic acids being the major cause of viral inactivation^12,13^. As most RNA extracted from coronavirus disease 2019 (COVID-19) patient samples are typically on the scale of under a few hundreds of thousands of copies roughly equating to a few picograms of RNA in the most concentrated of samples, reverse transcription quantitative polymerase chain reaction (RT-qPCR) and reverse transcription loop-mediated amplification (RT-LAMP) have shown to be the primary assays for sensitive diagnostic detection on the scale of accurately and reliably detecting SARS-CoV-2, even down to 10s of copies of virus per sample. While these methods are reliable for diagnostic detection, they do not reflect inactivation or infectivity of virus. The gold standard for determining viral infectivity has traditionally been the viral plaque assay; however, this method requires handling of infectious material typically in a BSL3 lab and above. As the 2019-nCoV pandemic continues to spread it then becomes necessary to develop a safe and practical method for labs not equipped with higher biosafety level cabinets to take part in COVID-19 research.

Here we demonstrated viral RNA degradation using Twist Synthetic SARS-CoV-2 RNA Control 2 (MN908947.3) and RNA from a protein coated and lipid bilayer enclosed non-replicative recombinant virus that closely resembles the wild type mammalian pathogenic virus via the AccuPlex™ SARS-CoV-2 Verification Panel. We used a modified Luna^®^ Universal Probe One-Step RT-qPCR Kit with the WarmStart RT step omitted and instead performed qPCR on cDNA synthesized from random hexamers and a Superscript IV polymerase in order to ensure conservative estimates of RNA degradation as cDNA synthesized in this manner is more robust against partial degradation. While the traditional gel electrophoresis approach to measure RNA quality requires larger amounts of RNA, RT-qPCR can be used as an alternative method to measure degradation as more intact gene transcripts in greater abundance become amplified sooner as compared to degraded RNA^14^.

## Results

We tested four distinct bacterial species commonly found in the hospital environment (*Escherichia coli* strain K12, *Enterococcus faecalis*, *Bacillus subtlis*, and *Deinococcus radiodurans*) for their susceptibility to ozone. These bacteria were grown and tested in triplicate on four high-traffic surfaces from the hospital equipment: catheters, blankets, hospital remote controls, and syringes, with positive and negative controls also included for comparison. All samples were treated with the same doze of ozone, at increasing lengths of time (30, 60, and 120 minutes), and CFU counts were compared between ozone-treated and controls. Control experiments consisted of allowing bacteria to grow without ozone treatment and then collecting samples as outlined in the **Methods** for each of the time points. For each surface and species tested colony forming units (CFUs) were counted, CFU/mL calculated, and the triplicates were averaged with standard error of the mean calculated to obtain ozone kill curves (**Figure 1**). These results showed as much as a three-log-fold changes in CFUs, with the majority of the impact observed within the first 30 minutes of treatment. These data showed that the ozone treatment causes significant reductions in all bacterial species tested across the various surfaces tested (p<0.001, Fisher’s Exact Test), with increasing degrees of efficacy as a function of time.

**Figure 1.**
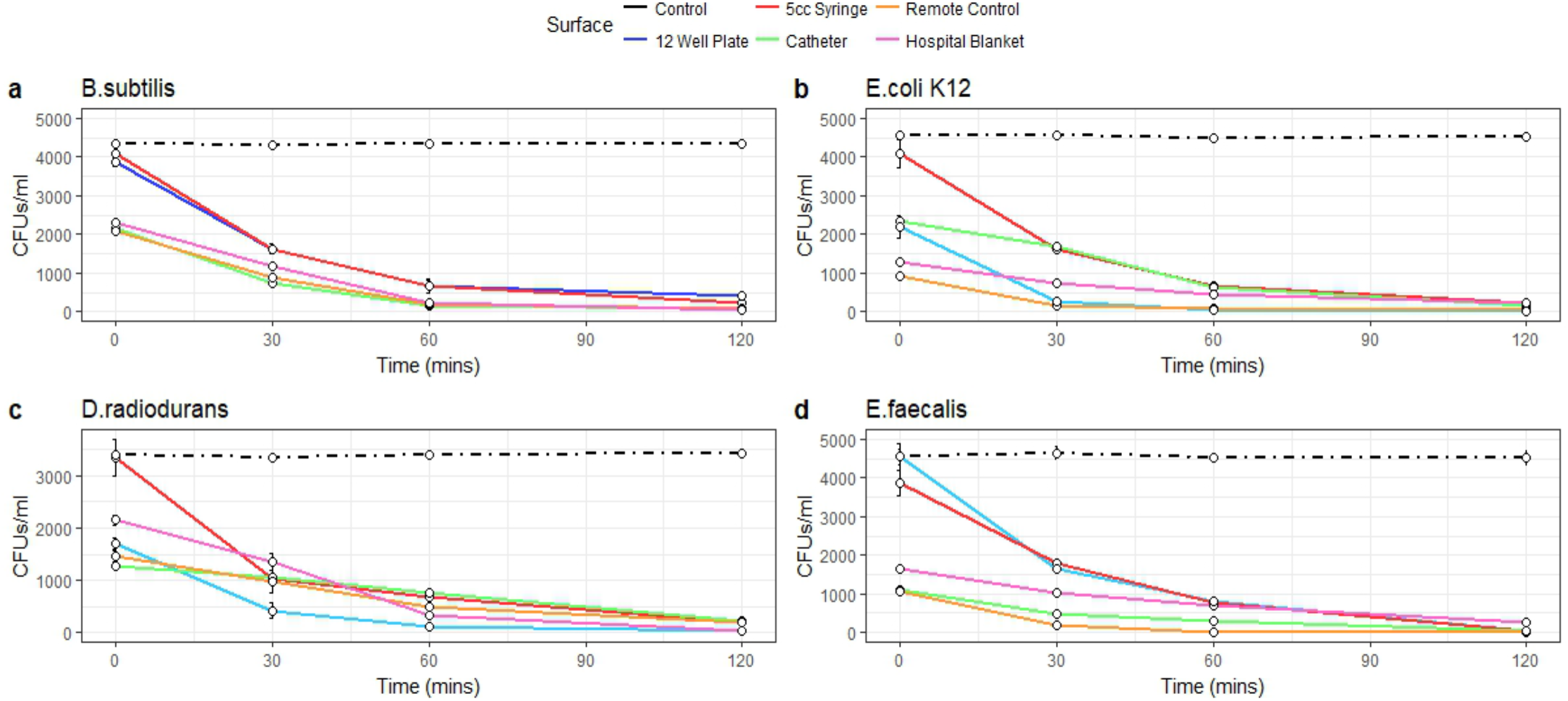
Ozone Kill Curves indicate ozone treatment reduces bacterial load. 1ml of bacteria for taxa *B. substilis* (a), *E. coli K12* (b), *D. radiodurans* (c), and *E. faecalis* (d) were pipetted into wells of 12 well plates and 100ul was collected at each time point for plating. 100ul of bacteria were pipetted inside 5cc syringes, inside catheters, onto remotes, and onto hospital blankets in triplicates and swabbed for 3 minutes at each time point for plating. Colony forming units were then counted following plating and 12hr incubation. CFUs are converted into CFUs/ml (y-axis). Error bars represent standard error of the mean. Controls are indicated as black lines. Ozone treatment (x-axis) is at 20ppm.

Moreover, both the different surfaces and different species showed distinct rates of reduction and response to the treatment. For example, *E. coli* showed the greatest sensitivity to killing in the plate, but less so on the remote controls. Also, the surface with the greatest impact (the most species killed) was the syringe, specifically at the 2-hour time point (>99.3%), vs. the greater variance on the hospital blankets. Also, the 2-hour time points for 12-well plate experiments showed a wide range of bacterial reduction across the different species, from 89.75% to 99.70%. The syringe experiments were the most consistent, with a range of bacterial reduction from 94.24% to 99.57%; conversely, the other surfaces showed a wider range of bacterial reduction, from 83.59% to 96.00% for the catheter, 86.10% to 99.31% for the remote control, and 82.99% to 98.36% for the hospital blankets. In all cases, these reductions were highlight significant (p<10^-5^) compared to the negative controls (0.0-0.4%).

Synthetic SARS-CoV-2 RNA was shown to have significant degradation 1hr into ozone treatment (p<0.001) using one way Anova Analysis and Tukey’s method (global p< 2.2e-16) among comparisons of cycle threshold (Ct) values obtained from our degradation robust modified qPCR method (**Supplemental Table 1)**. While some non-significant degradation of RNA occurred in non-treated controls left to air exposure on a bench top, degradation was significant for samples 1hr with percent RNA log fold change of 4hr time points nearing complete RNA degradation as compared to RNAse treated samples. Control samples treated with air exhibited an RNA Fold change of 98.69%, 93.55%, 86.22%, 61.78%, and 50.42% of intact amplifiable RNA remaining for 30mins, 1hr, 2hr, 3hr, and 4hrs respectively with 0hr control representing 100% of RNA; while ozone treated samples exhibited 65.13%, 25.82%, 11.24%, 12.46% and 6.16% of RNA remaining for 30mins, 1hr, 2hr, 3hr, and 4hrs. RNAse treated samples represents complete degradation with 0% intact amplifiable RNA remaining. Error bars represent variance (**Figure 3a)**.

**Figure 2.**
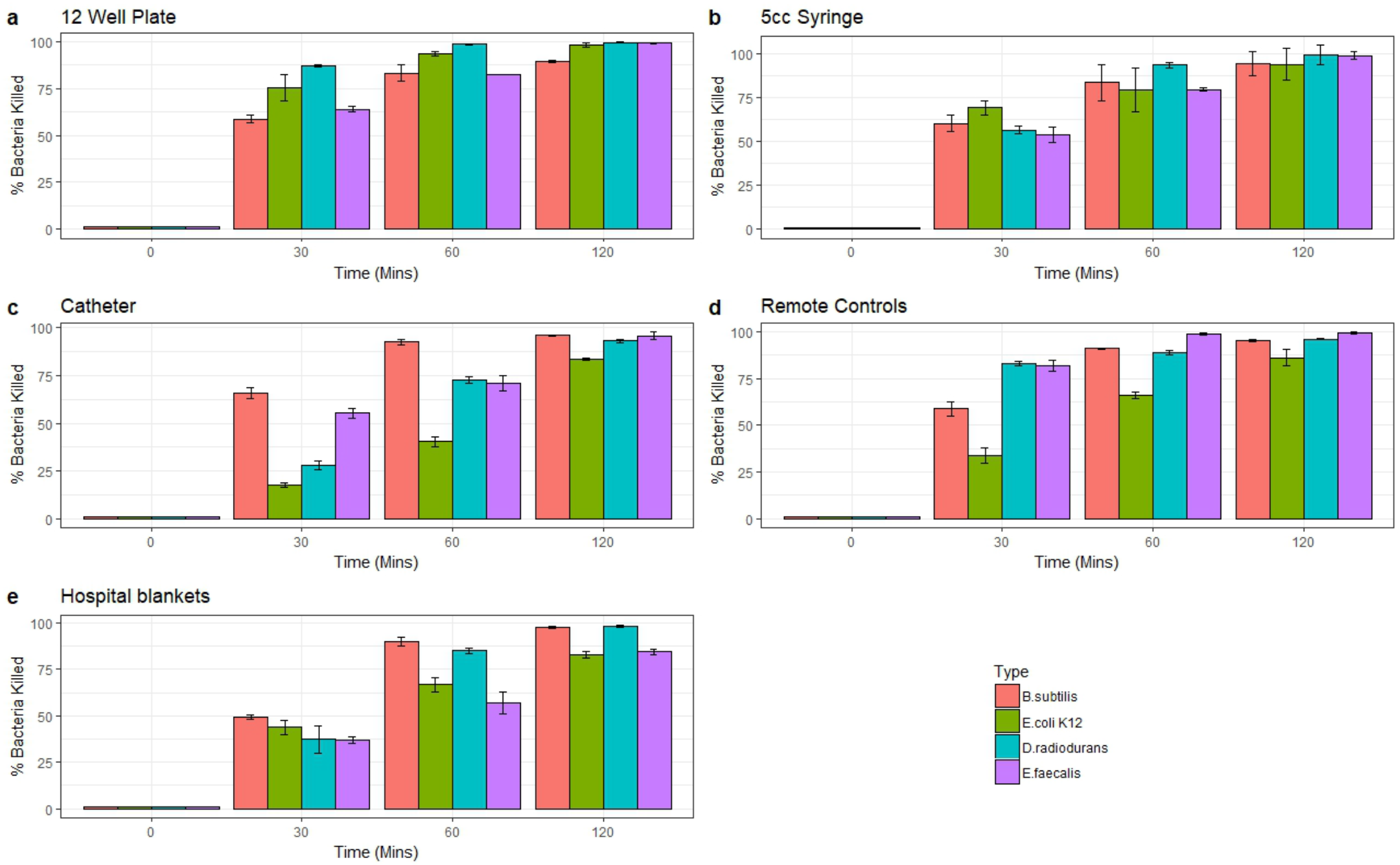
Percent Bacterial Reduction. Bacterial percent reductions for *B. substilis* (red), *E. coli K12* (green), *D. radiodurans* (blue), and *E. faecalis* (purple) from ozone treatment (20ppm). Error bars represent standard error of the mean from triplicate experiments on a variety of surfaces: a) 12 well plates b) syringes c) catheters d) remote controls e) hospital blankets. All differences compared to control are statistically significant (p<0.001, Fisher’s Exact Test).

**Figure 3.**
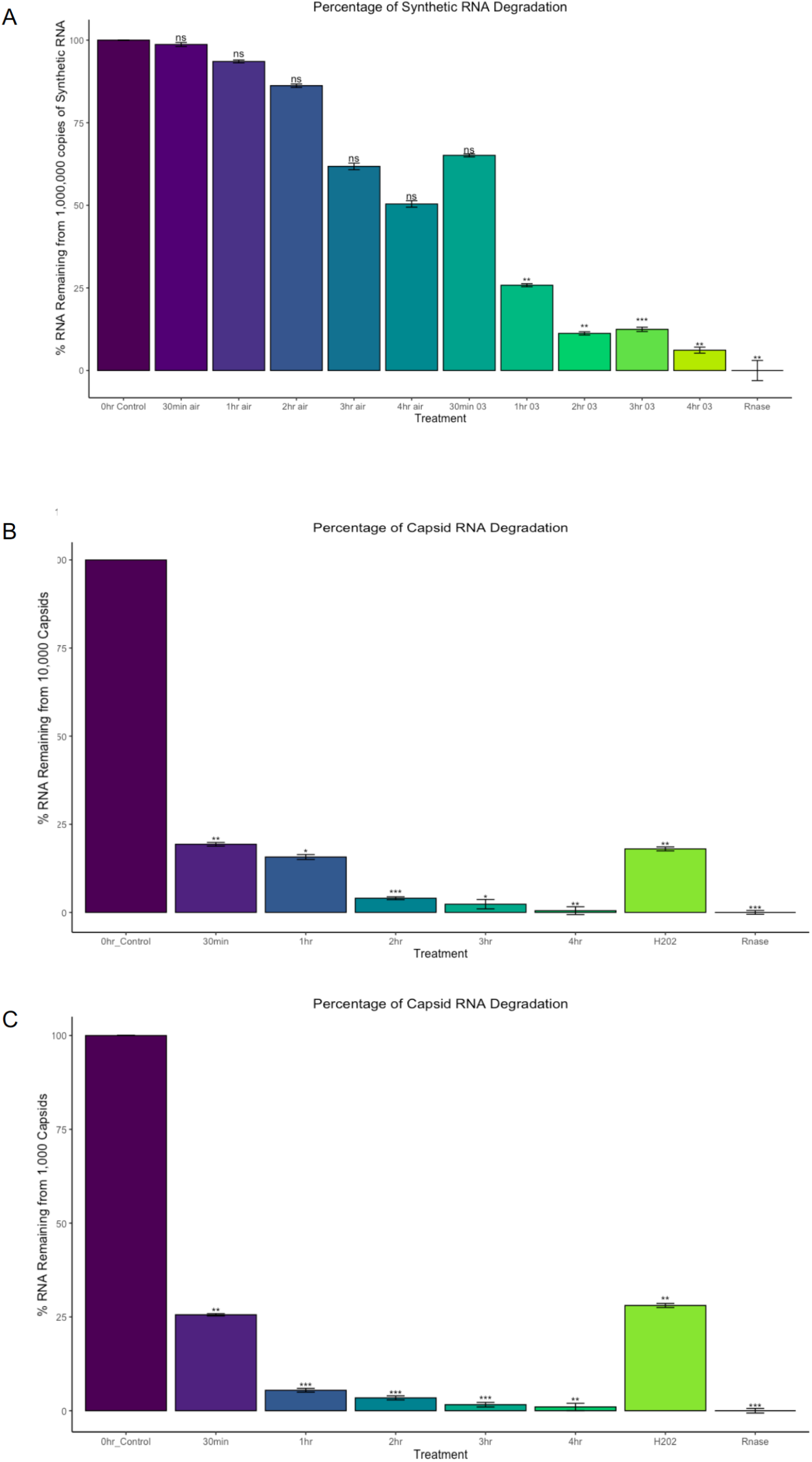
SARS-CoV-2 percent log fold change between ozone treatments and controls. (A) 1,000,0000 copies of Synthetic SARS-CoV-2. (B) 10,000 copies of non-replicative SARS-CoV-2. Global p < 3.2e-13. (C) 1,000 copies of non-replicative SARS-CoV-2. 0hr no treatment control was used as the comparison reference for ozone treated samples and control samples left on the benchtop exposed to air. Rnase values indicates maximum RNA degradation. Treatment Ct values were first subtracted from 0hr control to obtain delta Ct values (ΔCT = CT target – CT reference). Delta Ct was then transformed into log scale using 2^-ΔCt^. Error bars represent variance and was calculated as s = (s12 + s22)^1/2, s= standard deviation. Global P-value were computed by one way Anova Analysis and Tukey’s method was used to distinguish P-values for multiple comparisons. Global p <2.2e-16. Significance codes 0***, 0.001**, 0.01*.

RNA from non-replicative capsid enclosed SARS-CoV-2 showed significant degradation 30mins into ozone treatment as could be reliably detected by 10,000 copies of virus (p<0.001) and 1,000 copies of virus (p<0.001) (**Supplemental Table 1**). 100uM of H202 treatment for 1 hour also produced significant degradation according to Ct values obtained compared to 0hr treatment (p<0.001 for both 10,000 and 1000 copies of virus) and RNA degradation was still comparable to 1hr of ozone treatment, p=.99. For 10,000 copies of virus the percent of log fold intact amplifiable treated RNA left over compared to 0hr control treatment representing 100% of RNA was 19.33%, 15.72%, 4.03%, 2.34%, 0.49% for 30mins, 1hr, 2hr, 3hr, and 4hrs of ozone treatment respectively. 1hr of H202 treated RNA left over was 18.02% and RNase treated samples represented completely degraded RNA with 0% RNA left (**Figure 3b**). 1,000 copies of virus showed similar results to the 10,000 copies of virus tested. The percent RNA log fold change of ozone treated samples compared to 0hrs was 25.58%, 5.46%, 3.43%, 1.61%, and 1.01% for 30mins, 1hr, 2hr, 3hr, and 4hrs of ozone treatment respectively. H_2_O_2_ treated samples were 28.06% degraded after and RNAse representing complete degradation was 0% (**Figure 3c**). Mean Ct values were obtained along with Ct standard deviation. To obtain changes in degradation delta Ct values were calculated as either treatment or timepoint mean Ct subtracted from 0min control mean Ct. From this RNA fold change vs. Control was calculated as 2^-ΔCt^ as the difference between one cycle reflects a 2 fold difference in starting transcript level^13,14^.

## Discussion

Antibiotic resistance is a major financial burden on the United States healthcare system, often associated with common infections in hospitals which are caused by the sharing of rooms of infected patients, transmission by hospital workers, overgrowth of pathogens in the patient’s own microbiome, and interactions with surfaces and equipment that harbor these pathogenic bacteria. Fortunately, alternatives to antibiotics are being developed to take on this looming challenge.

Our findings provide support that ozone treatment is an effective sterilization method to combat HAIs in medical environments. We report rapid killing of the majority of medically relevant bacteria within 30 minutes, and up to a 99% reduction in viable bacteria at the end of 2-hour treatment cycles, with as much as a 3-4-fold log kill range. Here, we used an ozone generator from Sani Sport, but without any modifications or changes to the instrument. As such, changes in the flow rate, pressure, or temperature could increase the efficacy of this method, and possibly reduce bacterial species on common hospital surfaces in less time (~10-15 minutes).

It is interesting to note that different surfaces led to variation in the reduction of the same bacterial species. For example, the syringe and catheter experiments killed off bacteria at a slower rate than remote control and 12 well plate experiments. A likely explanation for this is that bacteria were inoculated directly onto the surfaces of 12-well plates and remote controls and so ozone could interact with more bacteria without any obstruction. Regardless, ozone treatment still demonstrated bacterial reductions even when inoculated inside tube-like structures such as catheters and syringes. Furthermore, ozone may not damage sensitive equipment (vs. bleach), which can ensure continued use of sterilized equipment in medical environments. Further work could explore the mechanisms by which bacteria respond to this treatment. Ozone is thought to disrupt membrane integrity, and so monitoring viability in this context can provide for a more conservative methodology^5^. While utilizing CFUs as a method to assess bacterial reduction have been the gold standard in many kill curve experiments, this method is limited in being able to discriminate viable but not cultivable cells (VBNC), and other methods such as propidium monoazide (PMA)-qPCR or flow cytometry could also be used^8^. In the context of further testing more pathogenic bacteria known to enter VBNC states it would be worthwhile to use these methods in conjunction with colony forming unit counts.

It would also be interesting to utilize RNA sequencing to study the stress response to ozone treatment in various HAI bacterial species. While our methods here demonstrate that 20 ppm of ozone can severely reduce several bacterial populations, there is little known about the stress response genes activated in response to ozone. The utilization of these stress responses are typically mediated by global regulatory mechanisms which affect biochemical pathways leading to physiological changes that confer survival. We hypothesize that regulatory networks such as heat shock, membrane integrity, and DNA damage may be activated. Particularly, it is interesting to note that ozone kills *D. radiodurans*, as this species has been known to have the unique ability to reconstruct its fragmented genome in response to ionizing radiation.^15^ There are a number of theories of how *D. radiodurans* is able to survive such extreme stressors one of them being an unusual capacity to avoid radiation induced protein oxidation, thus it would be interesting to study the stress response pathways in the context of Sani Sport ozone mediated treatment^16–18^. Understanding these mechanisms can be valuable for discovering novel genes for a multitude of bacterial species to decipher phenotypic characteristics, virulence regulation, and survivability, which may impact other medical environments like ambulances^19^ and broader urban environments around the world^20^.

Here we describe a new method for determining RNA degradation in a partial degradation robust manner for SARS-CoV-2 utilizing replication-deficient non-infectious virus that closely resembles the wild-type virus. While RNA degradation has been shown to be the main mechanism for viral inactivation^12,13^ in order to further validate these results traditional viral plaque assays using infectious SARS-CoV-2 virus must be employed. Nonetheless, this method provides for a safe way for labs not equipped with higher biosafety level cabinets to measure efficacy of viricidal treatment. As it has been demonstrated that N95 mask filtration efficiency is resistant to damage via many cycles or high dosage of ozone^8,9^, ozone can be a viable option for sterilization of PPE but also many different types of surfaces as well. While our limit of detection using our primers and qPCR chemistry was 1000 copies of non-infectious virus; this methodology could also be adapted to other primer sets and qPCR kits to perhaps measure even lower amounts of copies of virus per sample^21^.

## Methods

### Ozone treatment of bacteria

We used four common HAI-related, bacterial species to treat with ozone: *Escherichia coli* strain K12 (substrain MG1655/ATCC 700926), *Enterococcus faecalis* (OG1RF ATCC 47077), *Bacillus subtlis* (subtilis strain 168 ATCC 23857), and one isolated strain *Deinococcus radiodurans* (R1). Single, isolated colonies were cultured and treated with ozone (20ppm) inside the continually-generated ozone instrument (Sani Sport Supreme) at 0, 30, 60, and 120 minutes, which represent common intervals for cleaning in medical settings. We inoculated various surfaces that are common in hospital settings and produced ozone kill curves for each surface and bacterial strain. After swabbing each surface in triplicate, we isolated the bacteria from the swabs and cultured them overnight on media agar plates from which we assessed bacterial death from counting colony forming units (CFUs). Plates were blinded during counting and annotated afterward.

From fresh grown stock solutions, 100ul of cultured bacteria were obtained and placed in 900ul of appropriate broth (tryptic soy broth or nutrient broth with 1% glucose) in 1.5ml eppendorf tubes. From this 1:10 dilution the bacteria were serially diluted to 1:100, 1:1000, 1:10,000, 1:100,000, 1:1,000,000, and 1:10,000,000. 100ul from each dilution was then plated on corresponding tryptic soy agar plates and nutrient broth with 1% glucose agar plates using Zymo glass shaker beads to randomly distribute colonies across the plate. Plates were then grown at 37°C overnight for 12 hrs. Plates that had about 300 CFUs were chosen for the Sani Sport ozone experiment to be able to distinguish between colonies.

Using the correct dilution, 1 ml of each single strain of bacteria was placed in triplicates in a 12-well plate, so as to have enough surface area exposed to ozone treatment. 3 samples were used for each time point of 0, 30, 60, and 120 minutes (min). Ozone treatment was done on separate days for each strain to reduce chance of cross contamination. The 12-well plate was placed uncovered inside of Sani Sport and for the 0 min time point, 100ul was taken from the 1 ml sample and placed in a labeled eppendorf tube to be plated at the end of the experiment. The remaining 900ul was placed in a cryotube with 70ul of DMSO added and then placed on dry ice before moving into −80°C freezer.

Sani Sport doors were closed and the “cycle” button was pushed. Each 16-minute cycle consists of a 10-minute phase where purified air is blown inside the machine from fans and followed by a 6-minute phase of 20ppm ozone treatment according to manufacturer’s instructions. Ozone is generated each cleaning cycle inside the machine by reacting oxygen available in the air with ultraviolet light. At the end of the 5^th^ cycle 100ul of sample was taken from the three corresponding 30min wells and saved for plating at the end of the experiment while 900ul is frozen down as described above. This is done for 5 more cycles for the 1 hour (h) time point and then 10 more cycles for the 2h time point. After the 2h time point the saved 100ul samples are plated on appropriate agar plates and labeled. CFUs are then counted the next day after a 12h incubation at 37°C.

The same methods are followed for the different surfaces tested (syringe, catheter, hospital blanket, etc.) with minor modifications. 100ul of bacteria sample were added to each surface via pipette in triplicates and then for each time point 0, 30, 60, and 120 minutes, the surface was swabbed for 3 mins using Isohelix buccal swabs. Bacteria were pipetted directly onto flat surfaces such as remote controls and hospital blankets while bacteria were pipette onto the inside surface of tube like structures such as catheters and 5cc syringes. For dry surfaces, swabs were wetted with respective media and then used for swabbing. Swabs were then placed in 1.5ml Eppendorf tubes and suspended in 100ul of corresponding media. After the 2h time point each Eppendorf tube containing swab and media was vortexed for 10 seconds at max speed to shake bacteria off of swab and into surrounding media at bottom of tube. The tubes were then centrifuged at 6000 rpm for 10 mins at room temperature to collect bacteria at bottom of tube. Swabs were removed using sterile tweezers and media was pipetted up and down 10 times to resuspend bacteria at the bottom of the tube. 100ul of media was then plated onto respective agar plates, labeled, and incubated at 37°C overnight for 12h. CFUs were counted the following day.

### Ozone Treatment of Synthetic SARS-CoV-2 RNA

10 ul aliquots of Twist Synthetic SARS-CoV-2 RNA Control 2 (MN908947.3) (1,000,000 copies/ul, 11pg/ul) were subjected to ozone treatments at 30 minutes (5 cycles), 1hr (10 cycles), 2 hrs (20 cycles), 3 hrs (30 cycles), and 4hrs (40 cycles) in open capped PCR RNase free PCR tubes in the Sani Sport Supreme. A 0min control that was immediately processed for first strand cDNA synthesis after thawing on ice from −80°C was used as the baseline Ct value from which no degradation occurred. Degradation controls were also included in the experiment in which 10uls of Twist Synthetic SARS-CoV-2 RNA was subjected to 1ul of 100mg/ml of RNAse A for 30mins at room temperature before being converted to cDNA. No treatment open air controls were also included to compare the degradation rate between normal bench top RNA degradation and degradation from ozone. All treatments and controls were performed in triplicates and cDNA synthesis was initiated at the end of each time point using Invitrogen SuperScript iV Reverse Transcriptase for sensitive and robust synthesis of partially degraded RNA.

Briefly, 1ul of 60uM of NEB random primer mix was annealed to 1ul of template RNA with 1ul of 10mM dNTP mix, and 10ul of nuclease free water at 65°C for 5 mins and then incubated on ice for 1 min. 4ul of 5x SSIV buffer, 1ul of 100mM DTT, 1ul of RNaseOUTä, and 1ul of SuperScriptÒ IV Reverse Transcriptase (200 U/ul) were combined with annealed RNA and the combined reaction mixture was incubated at 23°C for 10mins, followed by 55°C for 10mins, and lastly the polymerase was heat inactivated by incubating at 80°C for 10 mins.

cDNA was then quantified using Qubit RNA HS Assay kit on a Qubit fluorometer to obtain cDNA concentrations of roughly 0.5ng/ul. 5ul of cDNA was then added to Luna^®^ Universal Probe One-Step Reaction Mix supplied at a 2X concentration containing Hot-Start Taq DNA Polymerase, dNTPs, and buffer for qPCR on a QuantStudio 6 Flex Real Time PCR system. 1.5ul each of a stock 22.5 nmol CDC 2019-nCoV-N1 Combined Primer/Probe Mix and 2019-nCoV_N2 Combined Primer/Probe Mix^22,23^ were added to the qPCR reaction with nuclease free water for a total reaction volume of 20ul in a 384 clear well PCR plate. Non template controls of nuclease free water were included to assess presence of contamination or primer dimers. A standard curve was also generated using 1:100 dilutions of stock Twist Synthetic SARS-CoV-2 RNA Control 2 for 10^7, 10^5, 10^3, and 10^1. All reactions done in triplicates. A standard Taqman Reaction protocol was set on the Quantstudio 6 with FAM reporter dye selected and no quencher. qPCR conditions consisted of an initial denaturation step of 95°C for 3 mins followed by 45 cycles of a 95°C denaturation step for 15 seconds and an extension step of 58°C for 30 seconds.

### Ozone Treatment of Non-Replicative Virus

To measure the effect of ozone on intact viral particles we subjected triplicates of non-replicative recombinant viruses via the AccuPlex™ SARS-CoV-2 verification panel to the same experimental conditions as our Twist Synthetic RNA experiment. 100ul of each 100,000 copies/ml, 10,000 copies/ml, and 1000 copies/ml were aliquoted into PCR tubes and subjected to ozone at 30mins, 1hr, 2h, 3hr, and 4hrs. Controls include a 0min time point and a 100uM H_2_O_2_ 1hr treatment. All three concentrations were used in order to establish the limits of detection for our modified RNA degradation qPCR assay.

At the end of each time point treatment RNA from non-replicative virus samples were obtained using Zymo Quick-RNA Viral kit according to the manufacturer’s instructions. Briefly, 100ul of 2x Concentrate DNA/RNA Shield™ was added directly to each sample. 400ul of Viral RNA Buffer was added to the mixture and then transferred to a Zymo-Spin™ IC Column in a collection tube and centrifuged for 2 minutes at 16,000 x g. The Column was then transferred to a new collection tube and 500ul of RNA Wash buffer was added to the column and centrifuged for 30 seconds at 16,000 x g twice followed by 500ul of a 95% ethanol wash step in which the column in a new collection tube was centrifuged for 1min at 16,000 x g. RNA was eluted in 6ul of nuclease free water and then was immediately subjected to cDNA synthesis and qPCR as mentioned above.

## Ethics Approval

None required for this study

## Contributions and Consent for Publication

CW, SR, DD, NBO, RO, DB, EA, CEM wrote and reviewed the text and data. CW performed the experiments with SR. RO, NBO, and CEM reviewed the data. All authors reviewed and approved of the manuscript.

## Availability of Data and Material

All raw data and CFU counts are included in the manuscript.

## Acknowledgements and Funding

We would like to thank funding from the Irma T. Hirschl and Monique Weill-Caulier Charitable Trusts, Bert L and N Kuggie Vallee Foundation, the WorldQuant Foundation, The Pershing Square Sohn Cancer Research Alliance as well as NASA (NNX14AH50G, NNX17AB26G), the National Institutes of Health (R25EB020393, R01NS076465, R01AI125416, R01ES021006, 1R21AI129851, 1R01MH117406), the Bill and Melinda Gates Foundation (OPP1151054).

